# Genetic inactivation of essential *HSF1* reveals an isolated transcriptional stress response selectively induced by protein misfolding

**DOI:** 10.1101/2023.05.05.539545

**Authors:** Michela Ciccarelli, Anna E. Masser, Jayasankar Mohanakrishnan Kaimal, Jordi Planells, Claes Andréasson

**Affiliations:** Department of Molecular Biosciences, The Wenner-Gren Institute, Stockholm University, S-10691 Stockholm, Sweden

**Keywords:** Proteostasis, heat shock factor 1, environmental stress response, heat shock proteins, heat shock, azetidine-2-carboxylic acid

## Abstract

Heat Shock Factor 1 (Hsf1) in yeast drives the basal transcription of key proteostasis factors and its activity is induced as part of the core heat shock response. Exploring Hsf1 specific functions has been challenging due to the essential nature of the *HSF1* gene and the extensive overlap of target promoters with environmental stress response (ESR) transcription factors Msn2 and Msn4 (Msn2/4). In this study, we constructed a viable *hsf1*Δ strain by replacing the *HSF1* open reading frame with genes that constitutively express Hsp40, Hsp70 and Hsp90 from Hsf1-independent promoters. Phenotypic analysis showed that the *hsf1*Δ strain grow slowly, is sensitive to heat as well as protein misfolding and accumulates protein aggregates. Transcriptome analysis revealed that the transcriptional response to protein misfolding induced by azetidine-2-carboxylic acid is fully dependent of Hsf1. In contrast, the *hsf1*Δ strain responded to heat shock through the ESR. Following HS, Hsf1 and Msn2/4 showed functional compensatory induction with stronger activation of the remaining stress pathway when the other branch was inactivated. Thus, we provide a long overdue genetic test of the function of Hsf1 in yeast using the novel *hsf1*Δ construct. Our data highlight that the accumulation of misfolded proteins is uniquely sensed by Hsf1-Hsp70 chaperone titration inducing a highly selective transcriptional stress response.

## Introduction

Heat Shock Factor 1 (Hsf1) is a highly conserved transcription factor in eukaryotes that in response to stress induces a protective transcriptional program called the heat-shock response (HSR) (Ritossa, 1996; Kim *et al*., 2013; Wang *et al*., 2013; Steurer *et al*., 2018). In *Saccharomyces cerevisiae*, a single and essential gene, *HSF1,* encodes Hsf1 which is a nuclear protein that functions as a trimeric transcription factor under both basal and stress-inducing conditions (Morano *et al*., 2012; Verghese *et al*., 2012). Hsf1 binds conserved sequences consisting of three inverted repeats of nGAAn called Heat Shock Elements (HSEs) located in the promoters of its target genes (Sorger and Pelham, 1987a; Bonner *et al*., 1994; Santoro *et al*., 1998; Sakurai and Takemori, 2007). Binding to HSE promoters varies between targets and under basal and stress conditions and around 43 genes are expressed under basal conditions. Under stress conditions, binding to the target promoters increases rapidly, leading to higher expression and induction of the core HSR involving classical heat-shock proteins (HSPs) (Pincus *et al*., 2018). The amplitude and stress responsiveness of individual genes are controlled by varying enrichment of Hsf1 at the different HSEs among the target genes (Li *et al*., 2006; Krakowiak *et al*., 2018). Thus, Hsf1 is responsible for the basal and stress-induced gene expression of heat shock proteins, activating them to different extents (Sorger and Pelham, 1987b; Jakobsen and Pelham, 1988; Wiederrecht *et al*., 1988; McDaniel *et al*., 1989; Gross *et al*., 1990, 1993; Morano *et al*., 1999; Hahn *et al*., 2004).

The sensing mechanism that Hsf1 employs to monitor stress damage in the cell has been under investigation for three decades, yet experimental data have only recently provided convincing evidence for the regulation of Hsf1 activity by Hsp70 chaperone titration (Bonner *et al*., 2000; Voellmy and Boellmann, 2007; Zheng *et al*., 2016; Krakowiak *et al*., 2018; Masser *et al*., 2019). Accordingly, available Hsp70 in the nucleus form a complex with Hsf1 and hinders the transcription factor from binding HSEs. Stress accelerates protein misfolding, particularly of newly translated proteins, and therefor decreases the levels of available Hsp70 by occupying its substrate-binding domain. As a consequence, Hsf1 is liberated from Hsp70 association, binds HSEs and induces the HSR. The stress induced system is returned to its resting state when Hsf1-driven expression of Hsp70 has restored chaperone availability (Masser *et al*., 2020)

Hsf1 is not the only transcription factor activated in response to stress. A plethora of stress conditions including glucose starvation, osmotic and heat stress, activates the two paralogous transcription factors Msn2 and Msn4 (Msn2/4) and together they are responsible for the regulation of the environmental stress response (ESR) (Martínez-Pastor *et al*., 1996; Schmitt and McEntee, 1996; Treger *et al*., 1998; Boy-Marcotte *et al*., 1999; Gasch *et al*., 2000). Under non-induced conditions, Msn2/4 are retained in an inactive state in the cytoplasm (Görner *et al*., 1998; Beck and Hall, 1999; De Wever *et al*., 2005). Stress induces their rapid nuclear accumulation where they bind Stress Response Elements (STREs) in the promoters target genes (Peter Lee et al., 2008) Many of the Msn2/4 target genes are under parallel transcriptional control by Hsf1 and hence carry both STREs and HSEs in their promoters, making the contribution of each pathway to their basal and induced expression hard to distinguish. The fact that *HSF1* is an essential gene has also made the genetic analysis difficult. Nevertheless, transient inactivation of Hsf1 by anchor-away technology has resulted in the identification of 18 genes that appear to be Hsf1-specific targets, including *SSA1* and *SSA2* (Hsp70), *HSC82* and *HSP82* (Hsp90), *FES1* (Hsp70 nucleotide exchange factor), *YDJ1* (Hsp70 J-domain protein) and *HSP104* (disaggregase) (Solís *et al*., 2016). Moreover, among these 18 genes, the dependency on Hsf1 for the basal expression of two of these genes, *SSA2* and *HSC82,* was proposed to explain why *HSF1* is essential based on their ability to restore growth under Hsf1-ablating conditions using the anchor away method (Solís *et al*., 2016). Thus, the genes uniquely activated by Hsf1 likely represents the essential core chaperone components of the HSR.

In this study we stringently test the role of Hsf1 under basal and stress conditions using a newly constructed *hsf1*Δ strain. We inactivated the essential *HSF1* by replacing it with gene cassettes that express *SSA2*, *HSC82* and *YDJ1* from constitutive promoters. Using this strain and comparing it with a *msn2/4*11 strain (Caballero *et al*., 2011), we clarify the role of Hsf1 under basal and stress conditions. Transcriptional profiles show that Hsf1 was solely responsible for responding to specific protein misfolding stress induced by the proline analogue azetidine-2-carboxylic acid (AzC). Hence, in cells Hsf1-Hsp70 appears to provide the sole mechanism that is sensitive enough to detect the damage and mount a transcriptional response when protein misfolding is induced. In contrast, during heat shock the parallel HSR (Hsf1) and ESR (Msn2/4) pathways function in a compensatory manner, resulting in increased activation of the remaining branch when the other one was inactivated. Overall, by analyzing the novel *hsf1*Δ strain, this study provides insight into the specificity of the transcriptional stress responses in yeast and how the Hsf1 and Msn2/4 branches are wired. The data highlight a critical role of Hsp70-titration to induce the activity of Hsf1 in response to the build-up of misfolded proteins.

## Results

### Genetic inactivation of *HSF1* results in temperature-sensitive growth, severe protein aggregation but does not block protein disaggregation

*HSF1* is an essential gene in yeast but the non-growth phenotype, associated to its loss, has been proposed to be suppressed by restoring the basal expression of Hsp70 and Hsp90 (Solís *et al*., 2016). This raised the possibility to investigate the role of Hsf1 by deleting its gene in the context of *SSA2* (*HSP70*) and *HSC82* (*HSP90)* expressed from constitutive promoters. We repeatedly failed to obtain viable *HSF1* knockouts using a cassette with *SSA2* under the *TDH3* promoter and *HSC82* under the *TEF1* promoter and reasoned that we also need a J-domain protein (JDP) expressed from a constitutive promoter to secure the functionality of the Hsp70 system. JDPs promote Hsp70 interaction with its substrates and induce hydrolysis of ATP (Kampinga *et al*., 2019). Accordingly, we additionally incorporated *YDJ1* under the control of the constitutive *ADH1* promoter in the knock cassette construct and the *HSF1* locus was inactivated by deleting the entire ORF using single-step homologous in a diploid strain followed by sporulation and the isolation of a haploid clone (See Materials and Methods). The resulting *hsf1*Δ strain was complemented by re-introducing the functional *HSF1* gene in the *hsf1*Δ locus creating a *HSF1* rescue strain that still harbored the extra copies of *SSA2*, *HSC82* and *YDJ1* with constitutive promoters (Figure 1A).

**Figure 1.**
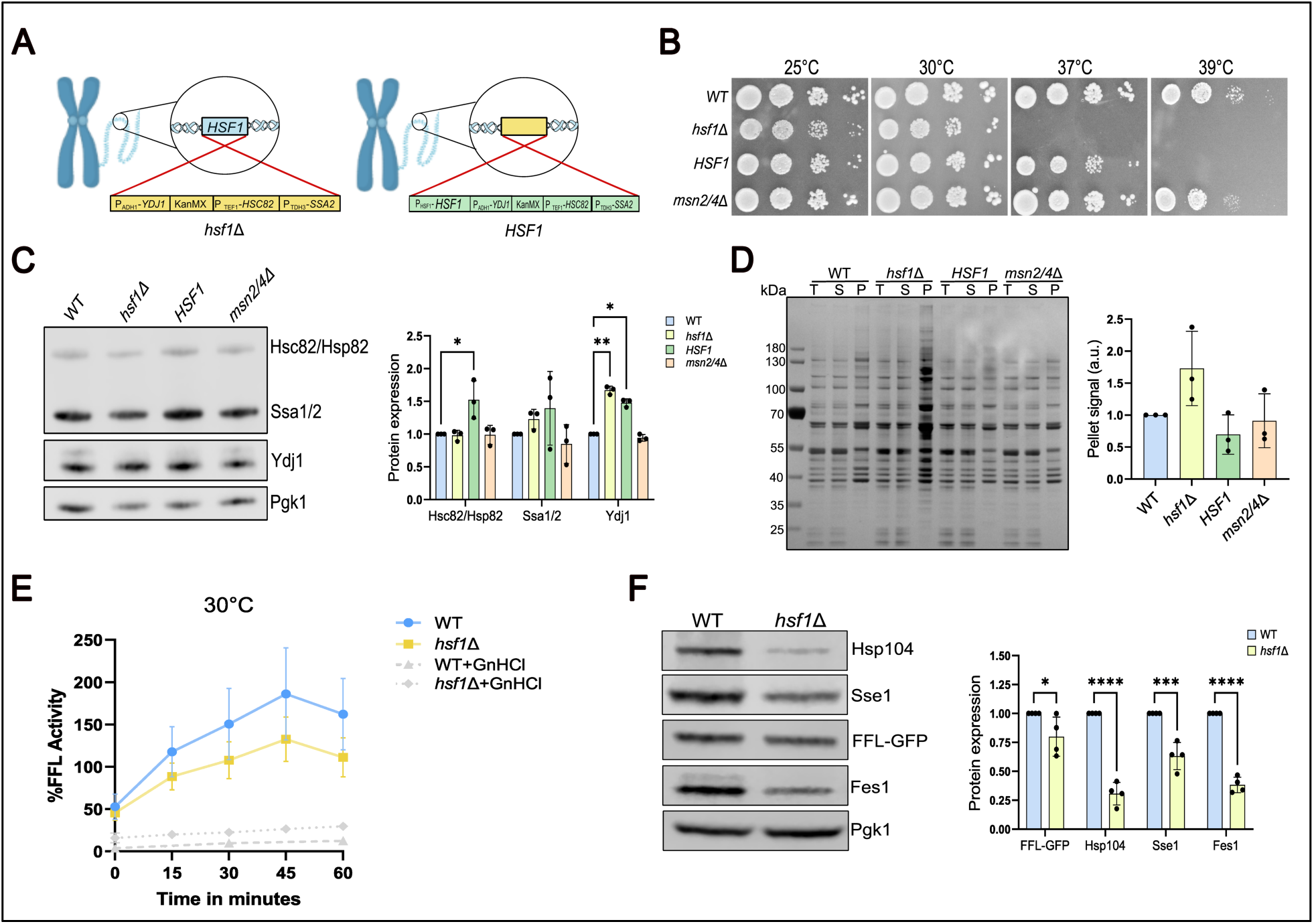
The *hsf1*Δ strain fails to grow at elevated temperatures and accumulates protein aggregates yet retains disaggregation activity. (**A**) The *hsf1*Δ and the *HSF1* rescue strain were constructed by integrating a gene cassette with *YDJ1*, *HSC82* and *SSA2* under constitutive promoters in the *HSF1* locus. The *HSF1* gene was then re-introduced in the *hsf1*Δ strain to create the rescued *HSF1* strain. (**B**) Growth characteristics of WT, *hsf1Δ, HSF1* and *msn2/4*Δ strains at 25°C, 30°C, 37°C and 39°C. Starting from the same cell density, 1:10 dilutions series were performed and cells were plated on SD medium supplemented with L-histidine, L-leucine, L-methionine and uracil and incubated for 3 days. (**C**) Left: western blot showing the protein levels of Hsc82/Hsp82, Ssa1/2 and Ydj1 at 30°C in all the strains. Right: western blot quantification; the protein signal was normalized to the respective Pgk1control signal. (**D**) Left: protein-aggregate isolate from of cells grown at 30°C. A silver stained gel shows total lysates (T), soluble fraction (S) and the aggregate pellet (P) of protein lysates from WT, *hsf1*Δ*, HSF1* and *msn2/4*Δ strains. Right: protein-aggregate quantification, the signal of the pellet was normalized to the respective signal of the total lysate. (**E**) FFL-GFP reactivation in the WT and *hsf1*Δ strains; cells carrying FFL-GFP were grown at 30° treated with CHX to stop translation (or CHX and 4 mM of GnHCl: grey dashed lines) and heat shocked at 43 °C for 15 minutes to inactivate FFL-GPF. FFL reactivation was measured 15, 30, 45 and 60 minutes after HS. (**F**) Left: western blot showing the protein levels of Hsp104, Sse1, FFL-GFP and Fes1 at 30°C in the WT and *hsf1*Δ. Right: western blot quantification; the protein signal was normalized on the respective control signal, Pgk1. Error bars show standard deviation of at least three independent experiments.

The growth of the constructed strains at the non-stressful temperatures (25°C and 30°C) and at more stressful temperatures 37°C and 39°C was assessed. A strain lacking *MSN2* and *MSN4* (*msn2/4*Δ) was also included in the analysis (Caballero *et al*., 2011). All the strains grew at 25°C and 30°C, however the growth of the *hsf1*Δ strain was significantly slower suggesting a severe proteostasis defect (Figure 1B). Increasing the temperature to 37°C and 39°C resulted in a complete failure of the *hsf1*Δ strain to form colonies and additionally unmasked a growth defect of the rescued *HSF1* strain. Overexpression of Hsp70 and Hsp90 is known to be associated with growth impairment and explains the phenotype of the complemented *HSF1* strain (Halladay and Craig, 1995; Keefer and True, 2017). Additionally, the *msn2/4*Δ strain displayed a modest growth impairment at 39°C. Next, we analyzed the protein expression levels of Ssa2, Hsc82 and Ydj1, the three HSPs introduced under constitutive promoters in the *hsf1*Δ and *HSF1* strains (Figure 1C). As predicted the Hsc82/Hsp82 levels were higher in the rescued *HSF1* while in the *hsf1*Δ stain the levels were comparable to the WT situation. Ssa1/2 levels followed a similar pattern but were slightly increased also in the *hsf1*Δ strain. The levels of Ydj1 were elevated in *hsf1*Δ and *HSF1* strains. The *msn2/4*Δ strain displayed no changes in the levels of any of the three chaperones. Consistent with that the activity of these transcription factors is low under logarithmic growth conditions. Thus, the *hsf1*Δ strain expresses Hsp70 and Hsp90 at levels comparable to wild type cells but exhibits severe growth impairment and the phenotype becomes more pronounced at elevated temperature stress conditions.

We considered the possibility that the *hsf1*Δ strain may display severely perturbed proteostasis already under basal conditions. To test our hypothesis, we isolated detergent resistant protein aggregates from cells grown at 30°C. Strikingly and in contrast to all the other strains, the *hsf1*Δ strain accumulated elevated levels of protein aggregates. Notably, despite extensively overlapping target promoters of Hsf1 and Msn2/4, the *msn2/4*Δ strain did not accumulate aggregates demonstrating that this is an Hsf1-dependent phenotype (Figure 1D). The rescued *HSF1* strain did also not accumulate aggregates, indicating that it is a specific consequence of Hsf1 inactivation and not the ectopic expression of *SSA2*, *HSC82* and *YDJ1.* Thus, despite the constitutive expression of Hsp70 and Hsp90, Hsf1 has an important role in maintaining proteostasis, suggesting a role for other factors that depend on Hsf1 for their basal expression.

We assessed whether the *hsf1*Δ strain maintained functional protein disaggregation and reactivation. These processes depend on JDPs, Hsp70, Hsp110 and Hsp104 that are all under transcriptional control of Hsf1 (Parsell *et al*., 1994; Glover and Lindquist, 1998; Kaimal *et al*., 2017; Sathyanarayanan *et al*., 2020). The WT and *hsf1*Δ strains carrying the reporter firefly luciferase (FFL) fused with GFP were grown at 30°C and treated with cycloheximide (CHX) to stop translation. FFL-GFP was inactivated by heat shock at 43°C for 15 minutes (Abrams and Morano, 2013; Kaimal *et al*., 2017). The *hsf1*Δ strain was able to reactivate the FFL activity although with slower kinetics than the WT strain (Figure 1E). The reactivation was as expected dependent on Hsp104, since it was completely blocked by the Hsp104 inhibitor guanidine hydrochloride (GnHCl) (Jung and Masison, 2001) (Figure 1E). The basal levels of Hsp104 were reduced in the *hsf1*Δ strain as were the levels of Hsp70 co-chaperones Sse1 and Fes1; this likely explains the slower kinetics of reactivation displayed by the knock strain (Figure 1F). Thus, the *hsf1*Δ strain maintains an overall functional protein disaggregation machinery.

### *The hsf1*Δ strain responds to heat shock via STRE elements

In order to test how the *hsf1*Δ strain responds to heat-shock stress, we heat-shocked cells at 37°C for 15 minutes and allowed them to grow out on solid medium at 30°C. The *hsf1*Δ strain as well as the other strains recovered from this insult without apparent loss of colony forming ability suggesting that this stress condition is useful to experimentally assessing the HSR (Figure 2A). To directly measure the activity of Hsf1 and Msn2/4 under heat shock conditions, we employed reporter plasmids that are based on NanoLuciferase under the control of a HSE or STRE dependent promoter, respectively (Hall *et al*., 2012; Masser A. et al., 2016; Masser *et al*., 2019). We pre-grew the strains at 25°C and subjected them to 15 minutes of HS at 37°C. The Hsf1-dependent HSR (HSE reporter) was readily detected in WT, *HSF1* and *msn2/4*Δ strains while, as predicted, it was not mounted in the *hsf1*Δ strain (Figure 2B, left). The Msn2/4-dependent ESR (STRE reporter) was in contrast functional in the *hsf1***Δ** strain supporting the notion that it does not depend on Hsf1 for stress signaling (Martínez-Pastor *et al*., 1996; Solís *et al*., 2016) (Figure 2B, right). The HSR and the ESR were suppressed in the Hsp70, Hsp90 and Ydj1 overexpressing *HSF1* rescue strain suggesting that these chaperones negatively impact not only on the induced activity of Hsf1 but also on Msn2/4. Notably the basal activity of Hsf1 was specifically suppressed in this strain, supporting the recent findings that Hsp70 directly binds to and attenuates the activity of Hsf1 (Zheng *et al*., 2016; Krakowiak *et al*., 2018; Masser *et al*., 2019).

**Figure 2.**
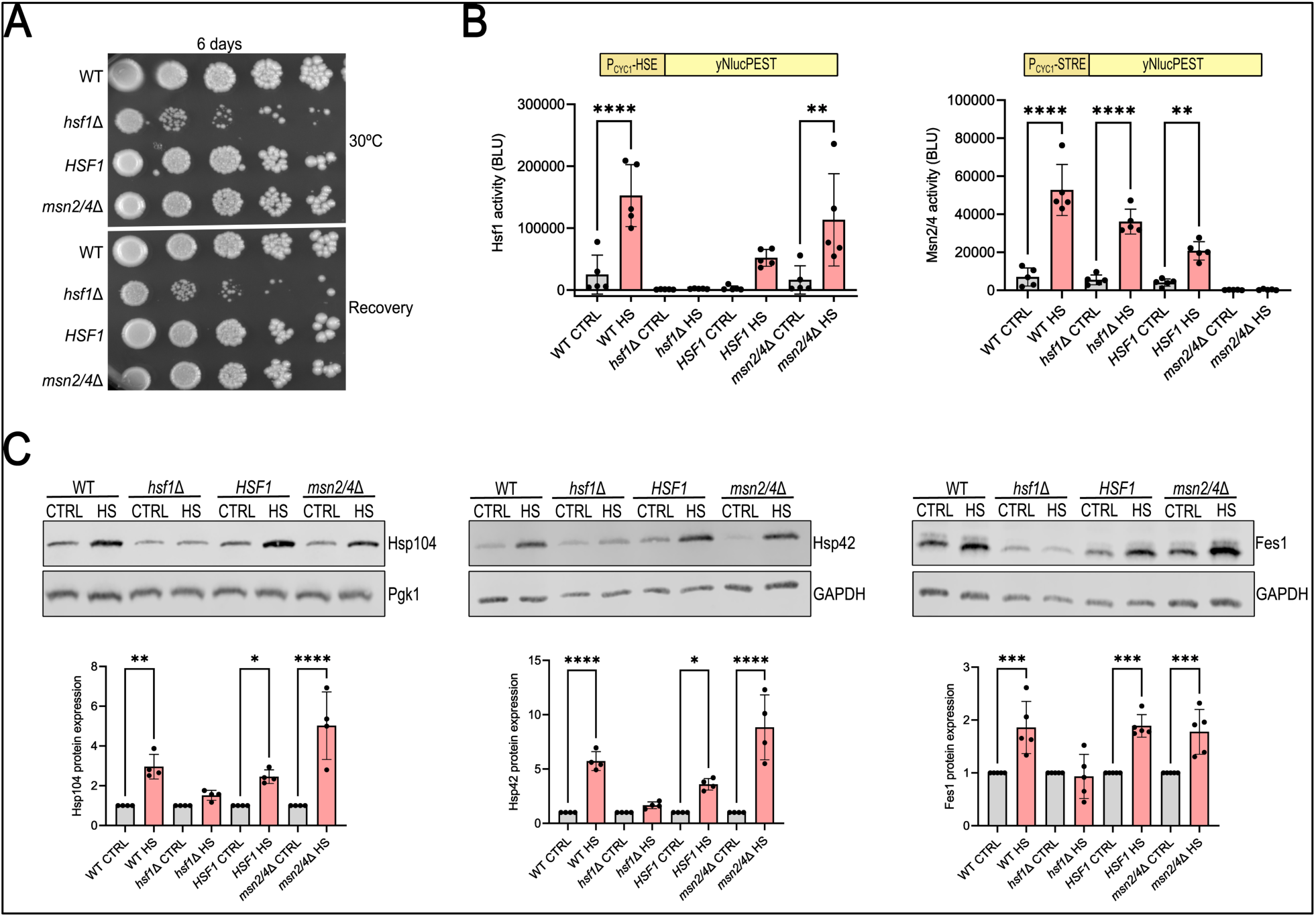
The *hsf1*Δ strain fails to induce the HSR following HS but maintains a functional ESR. (A) Growth assay of WT, *hsf1*Δ, *HSF1* and *msn2/4*Δ strains. Serial 1:10 dilutions were performed and cells were plated on SD medium or first heat-shocked at 37 °C for 15 minutes and then plated on SD medium (Recovery) and plates were incubated at 30°C for 6 days. (B) The activity of Hsf1 and Msn2/4 was measured using HSE and STRE NanoLuc reporters in the indicated strains before and after heat shock at 37°C for 15 minutes (grey: cells grown at 25 °C (CTRL), red: heat shock (HS)). (C) Top: western blot showing the protein levels of three Hsf1-selective targets: Hsp104, Hsp42 and Fes1 before and after HS at 37°C for 15 minutes in WT, *hsf1*Δ, *HSF1* and *msn2/4*Δ. Bottom: western blot quantification; the protein signal was normalized on the respective control signal (Pgk1 or GAPDH) (grey: cells grown at 25 °C (CTRL), red: heat shock (HS)). Error bars show standard deviation of at least three independent experiments.

Next, we assessed the HSR by Western blotting of Hsp104, Hsp42 and Fes1, three HSPs encoded by genes whose basal expression is selectively regulated by Hsf1 (Solís *et al*., 2016). The HSR induction of all three HSPs was largely attenuated in the *hsf1***Δ** strain, yet Hsp104 and Hsp42 levels did increase somewhat, likely as a consequence of that these proteins become part of stable protein aggregates following heat shock that delays their turnover, or that weak STRE sites are present in their promoters (Figure 2C) (Grably *et al*., 2002; Kuang *et al*., 2017) Overall, cells lacking Hsf1 can still respond transcriptionally to a HS via Msn2/4-dependent STRE promoters but fail to mount a transcriptional response via HSEs.

### *The hsf1*Δ strain is hypersensitive to the protein misfolding stressor AzC

HS not only impairs proteostasis but generally damages cells (Lambowitz et al. 1983; Patriarca and Maresca 1990; Toivola et al. 2010; Vogel, Parsell, and Lindquist 1995; Welch and Suhan 1985). In order to assess the response of the *hsf1*11 strain to a specific proteostasis insult, we treated cells with L-azetidine-2-carboxylic acid (AzC), a proline analogue that is incorporated into proteins when proline codons are translated and due to its 4-carbon ring impairs protein folding (Fowden, L., and Richmond, 1963; Zagari *et al*., 1990, 1994). Assessing a range of concentrations of AzC showed that WT, *HSF1* and *msn2/4*Δ strains displayed impaired growth at 10 mM AzC at 25°C and at 5 and 10 mM at 30°C (Figure 3A). In contrast, the *hsf1*11 strain was hypersensitive to AzC and displayed a growth defect that increased with increasing AzC concentrations at all temperatures. Protein aggregate analysis showed that 1 h treatment with 10 mM AzC resulted in substantially induced aggregation in both the WT and *hsf1*Δ strains (Figure 3B). Importantly, the *hsf1*11 strain accumulated aggregates already before the treatment but still AzC induced further protein aggregation, demonstrating that the strain received additional proteostasis damage upon AzC treatment. Thus, the *hsf1*Δ strain is hypersensitive to AzC and the compound rapidly induces protein aggregation in WT as well as in the *hsf1*11 strain.

**Figure 3.**
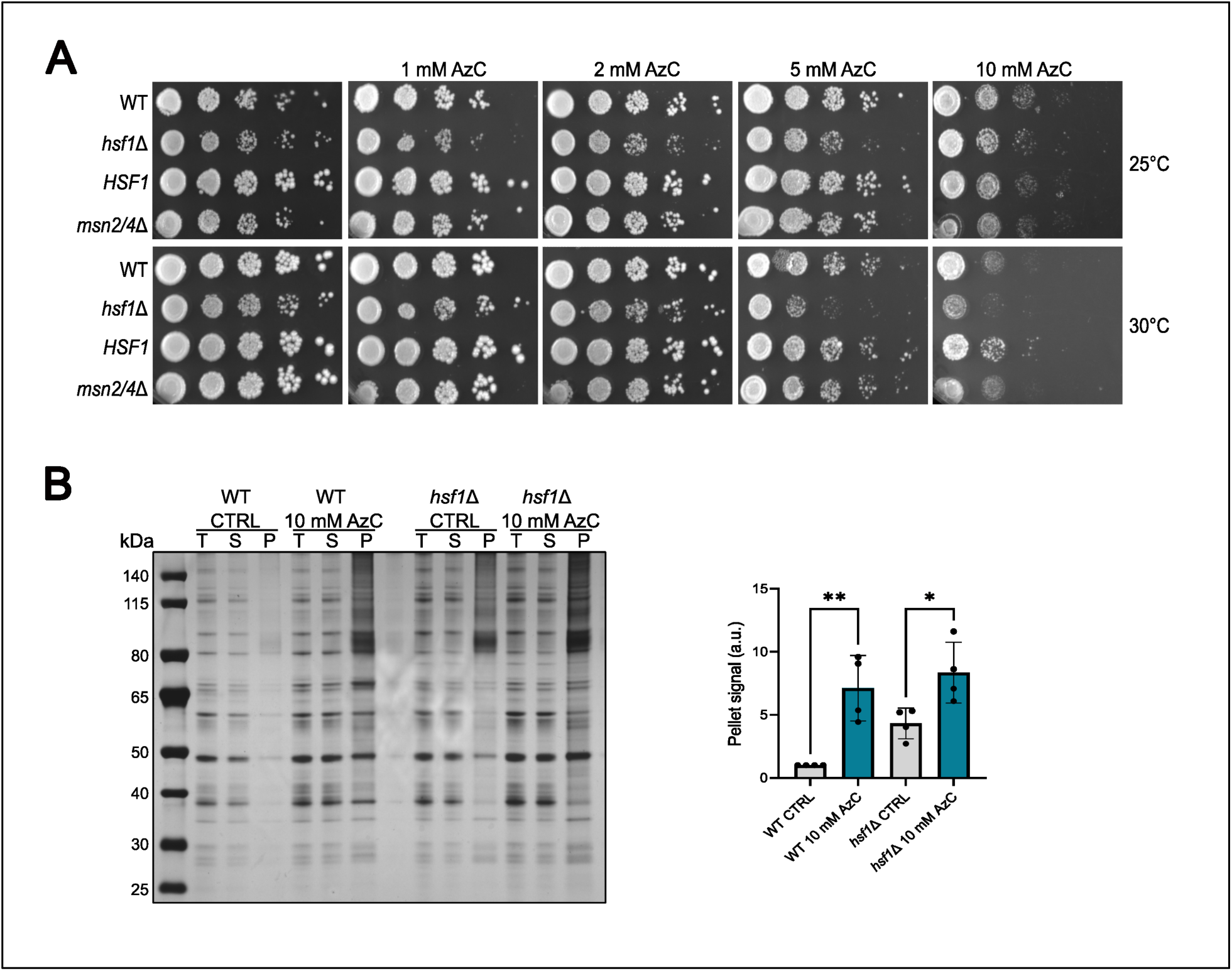
The *hsf1*Δ strain is hypersensitive to AzC and it induces widespread protein aggregation. (A) Growth assay of all the strains performed in YPD plates supplemented with 0, 1, 2, 5 or 10 mM of AzC at 25°C or 30°C. (B) Left: protein-aggregate analysis performed before (CTRL) and after treatment with 10 mM AzC for 1h at 25°C. Silver stain shows total lysates (T), soluble fraction (S) and the aggregate pellet (P) of protein lysates from WT and *hsf1*Δ. Right: protein-aggregate quantification; the pellet fraction was quantified and the signal of the pellet was normalized to the respective signal of the total lysate (grey: cells grown at 25 °C (CTRL); blue: 10 mM AzC). Error bars show standard deviation of at least three independent experiments.

### The transcriptional response to AzC-induced protein misfolding selectively depends on Hsf1

To assess how cells mount transcriptional stress responses when Hsf1 or Msn2/4 are inactivated, we performed mRNA-sequencing (mRNA-seq). The four strains WT, *hsf1*11, *HSF1* and *msn2/4*11 were grown at 25°C and treated with 10 mM AzC for 1 h or alternatively were heat shocked at 37°C for 15 minutes before RNA extraction and sequencing (Figure 4A). Principal component analysis (PCA) of the mRNA reads showed that all strains clustered under the control condition (growth at 25°C) and that the WT and *msn2/4*11 strains as well as the *hsf1*11 and *HSF1* strains paired up. Heat-shock resulted in large transcriptional changes and clustering of WT, *hsf1*11 and *HSF1* strains but not of the *msn2/4*11 strain (Figure 4B, left and middle panels). Around 2000 genes were differentially expressed after HS in all four strains and the overall response remained despite inactivation of Msn2/4 or Hsf1 (Figure 4C, Table 1). The data are consistent with previously published results showing that Hsf1 and Msn2/4 are responsible for subroutines in the larger HS stress transcriptional program and that the role of Msn2/4 is broader than that of Hsf1 (Solís *et al*., 2016). AzC treatment resulted in induction of a somewhat more compact transcriptional program involving nearly 1300 differentially expressed genes in wild type cells (Table 1). Importantly, the transcriptional response to AzC was nearly abolished in the *hsf1*11 strain. The rescued *HSF1* strain displayed an attenuated HSR upon AzC treatment (Figure 4B, left and right panels and Fig 4C, right panel, Table 1), recapitulating what we observed using the NanoLuciferase based Hsf1 reporters. Again, this illustrates how the overexpressed Hsp70 maintains Hsf1 negatively regulated by chaperone titration.

**Figure 4.**
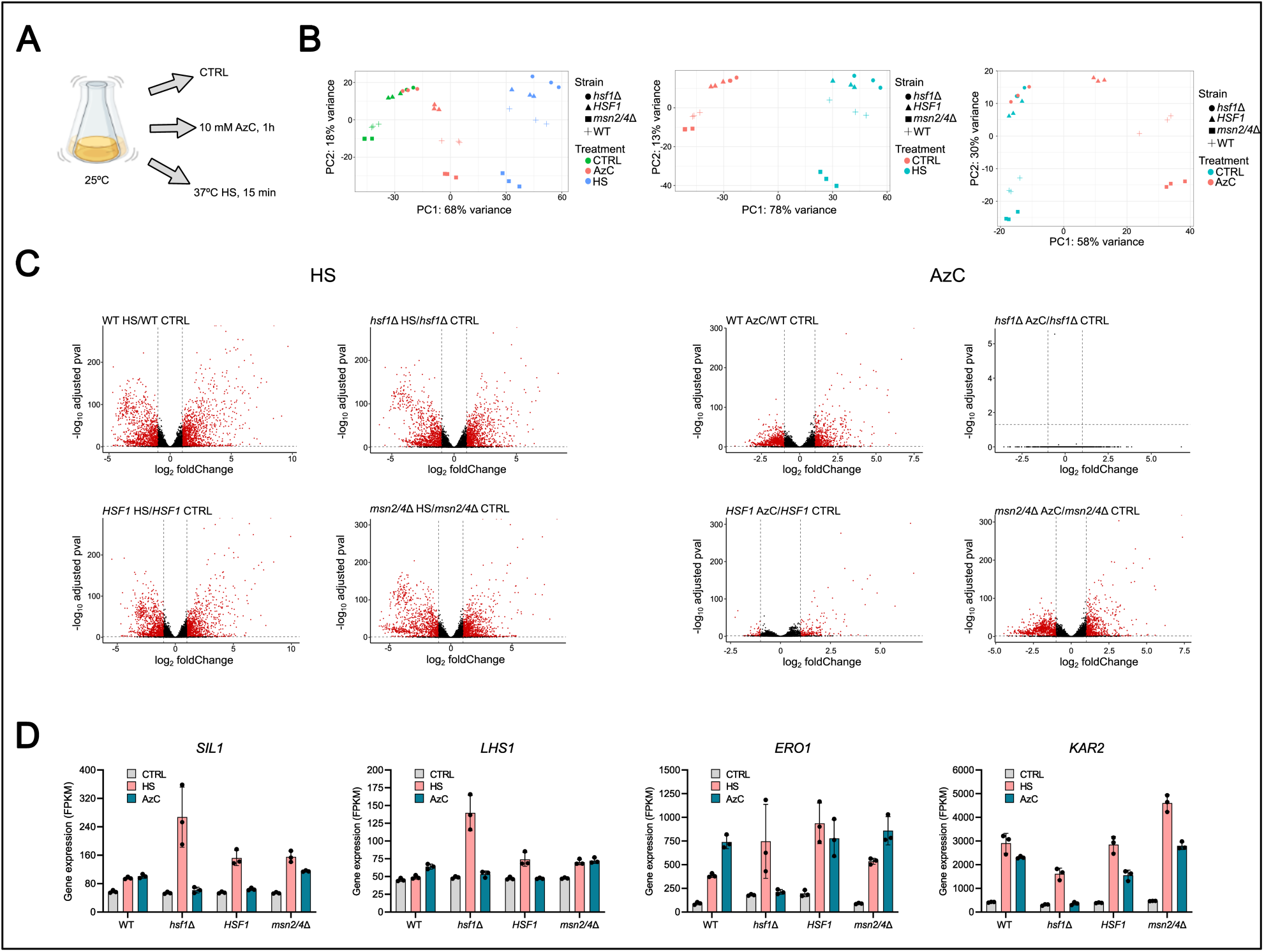
The HSR induced by protein misfolding (AzC) is selectively dependent of Hsf1 while HS induces both Msn2/4 and Hsf1. (A) Experimental workflow for transcriptome analysis by mRNA-seq. Cells were grown at 25°C (CTRL) and were subjected to HS at 37°C for 15 minutes or treated with 10 mM AzC for 1 h at 25°C. (B) Principal component (PC) analysis of transcriptome mRNA-seq data of all the strains at all conditions (left), HS treatment and CTRL (middle panel) or AzC treatment and CTRL (right). (C) Volcano plots showing gene expression changes after HS (left) and AzC treatment (right). Treated conditions are compared to CTRL conditions (cells grown at 25°C) for all the strains; genes with adjusted p value < 0.01 and absolute log_2_ Fold Change > 1 are considered significant. Threshold are indicated with dotted black lines. (D) Gene expression of 2 UPR-dependent targets (*SIL1*, *LHS1*) and 2 targets shared between Hac1 and Hsf1 (*ERO1* and *KAR2*) in all the strains in control conditions, upon HS or AzC treatment. Experiments were performed in triplicate.

**Table 1.** mRNAs abundance changes upon HS and AzC treatment

| Comparison | Number_DEG | Upregulated | Downregulated |
| --- | --- | --- | --- |
| WT HS/WT CTRL | 2232 | 1150 | 1082 |
| *hsf1*Δ HS/*hsf1*Δ CTRL | 1935 | 936 | 999 |
| *HSF1* HS/*HSF1* CTRL | 1816 | 862 | 954 |
| *msn2/4*Δ HS/*msn2/4*Δ CTRL | 1914 | 944 | 970 |
| WT AzC/WT CTRL | 1269 | 588 | 681 |
| *hsf1*Δ AzC/*hsf1*Δ CTRL | 0 | 0 | 0 |
| *HSF1* AzC/*HSF1* CTRL | 237 | 179 | 58 |
| *msn2/4*Δ AzC/*msn2/4*Δ CTRL | 1363 | 612 | 751 |

Number of differentially expressed genes (DEG) and up- and downregulated genes after heat shock at 37°C for 15 minutes (HS) or treated with 10 mM AzC for 1h at 25°C (AzC).

Hsf1 also activates promoters of secretory genes that are regulated by the Unfolded Protein Response (UPR) with its key transcription factor Hac1 (Ron & Walter, 2007). Hac1 expression is induced by unfolded/misfolded proteins in the ER using a chaperone-titration mechanism that involves negative regulation by secretory Hsp70 Kar2 (Cox and Walter, 1996; Chapman and Walter, 1997; Kawahara *et al*., 1997; Sidrauski and Walter, 1997) In the WT strain, UPR-responsive genes *LHS1* and *SIL1* responded weakly to HS and AzC, while *ERO1* and *KAR2*, that are known to be regulated by both UPR and HSR (Hahn *et al*., 2004; Yamamoto *et al*., 2005), responded robustly to the insults. The response to AzC was fully dependent on Hsf1 since it was not mounted in the *hsf1*11 strain. In contrast, the response to HS was much stronger when Hsf1 was inactivated (Figure 4D). Hence, Hsf1-Hsp70 and not the UPR senses protein misfolding induced by AzC, nevertheless Hsf1 has an important role in maintaining ER proteostasis leading to that its inactivation makes the UPR more responsive to HS damage (Liu and Chang, 2008). Overall, the transcriptional response to protein misfolding induced by AzC is fully dependent on Hsf1 activity and does not depend on the ESR or UPR, while all three pathways respond to HS.

Inspection of a group of genes that carry both HSE and STRE (Pincus *et al*., 2018) (e.g. *EDC2*, *GPH1*, *GRE3*, *SPI1*, *SSE2* and *TMA10*) revealed that they failed to respond to AzC treatment in the *hsf1*11 strain but remained responsive to HS (Figure 5A). The expression of previously defined Hsf1-selective target genes (Solís *et al*., 2016) were fully dependent on Hsf1 both for HS and AzC treatment (Figure 5B, top). Inspection of Msn2/4 target genes showed that their HS induction overall was dependent on Msn2/4 and not on Hsf1 (Figure 5B, bottom). In contrast, their induction in response to AzC treatment was fully dependent on Hsf1 indicating that only genes regulated by both Msn2/4 and Hsf1 responded to the AzC treatment. Furthermore, 18 reported Hsf1 selective target genes (Solís *et al*., 2016) were all induced upon either HS or AzC treatment in WT strain and their induction by AzC treatment was fully suppressed in *hsf1*Δ strain (Figure 5, C-D). Four of these 18 genes (*HSP82*, *HSP78*, *HSP42*, and *HSP104)* still responded to HS in the *hsf1*Δ strain and these genes have been reported to be regulated by Msn2/4 under stress (Grably *et al*., 2002; Kuang *et al*., 2017). Thus, AzC selectively induces Hsf1 activation and not Msn2/4 and cells are not able to transcriptionally respond to such protein folding damage in the absence of Hsf1.

**Figure 5.**
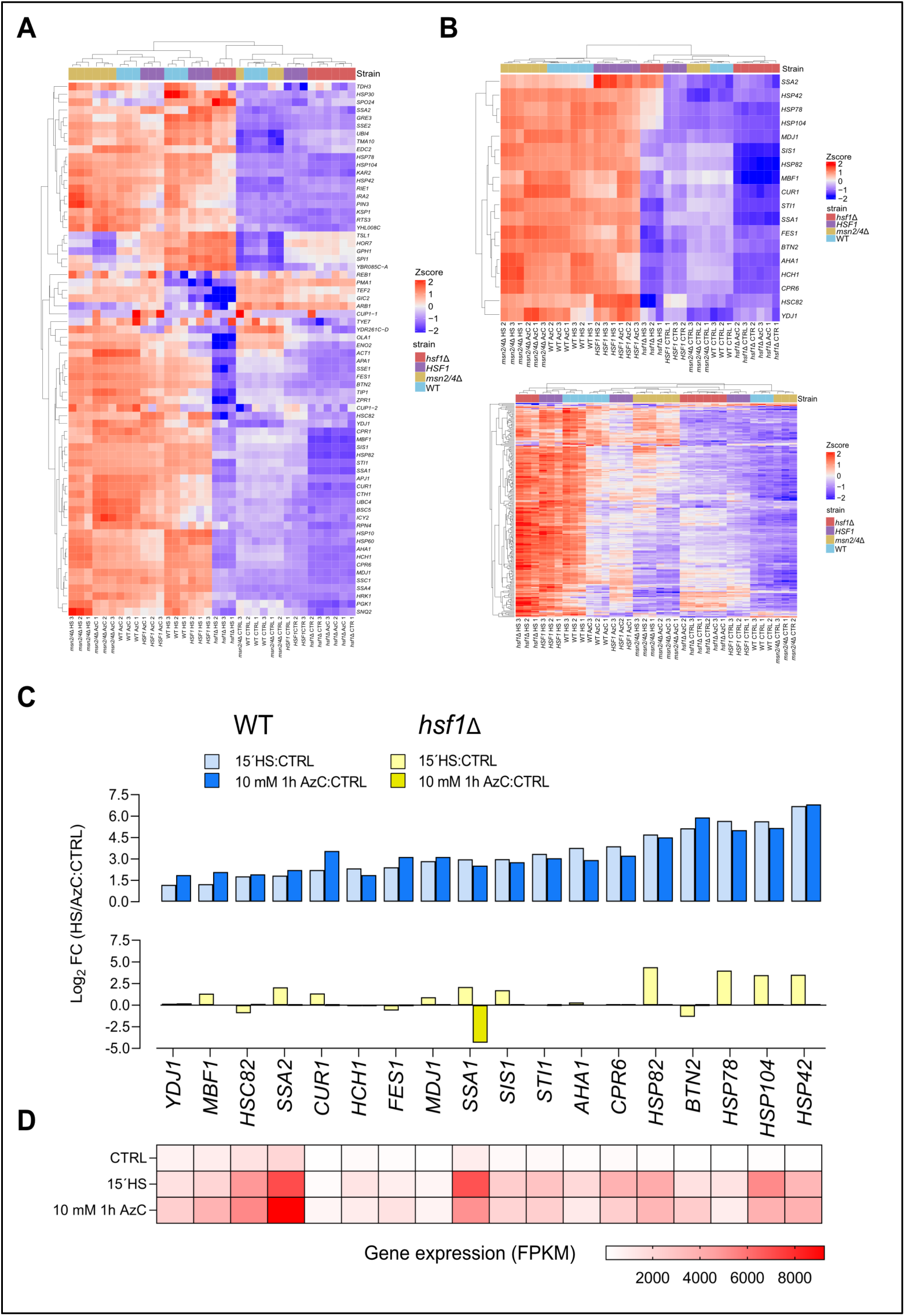
HS and AzC stress induce expression of Hsf1 and Msn2/4 target genes. (A) Heat map showing differential expression of all Hsf1 target genes in CTRL, HS and AzC treatment conditions in WT, *hsf1*Δ*, HSF1* and *msn2/4*Δ. (B) Heatmaps showing changes in the expression of Hsf1 selective targets (top) and Msn2/4 targets (bottom) in CTRL, HS and AzC treatment in all the strains. (C) RNA expression changes for 18 Hsf1 specific target genes in WT (top) and *hsf1*Δ (bottom) strains. (D) Heatmap showing changes in the expression of the corresponding 18 Hsf1 specific targets in the WT strain. Data from triplicates are shown.

### Hsf1 and Msn2/4 display functional compensatory induction following HS

Hsf1 and Msn2/4 function in parallel HS stress responsive pathways that in part converge on shared target promoters. This raises the question if inactivation of one of the pathways impacts on the activity of the other one. Indeed, the overall expression of Hsf1-selective target genes was found to be higher expressed in the *msn2/4*Δ strain than in the WT strain following heat shock (Figure 6A). Conversely, Msn2/4 target genes were found to be higher expressed in the *hsf1*Δ strain than in the WT strain following heat shock (Figure 6B). Next, we analyzed the expression of individual genes in the CTRL and HS conditions by selecting *FES1*, *BTN2* and *HCH1* as representatives of Hsf1 target genes and *CTT1*, *TPS3* and *TSL1* as representatives of Msn2/4 targets. In each case the induction was higher than in the WT strain when the parallel functioning transcription factor had been inactivated (Figure 6, C-D). Hence, cells mount transcriptional programs in response to HS that depend on both Hsf1 and Msn2/4 and when one pathway is inactivated, the remaining one responds stronger in a compensatory manner.

**Figure 6.**
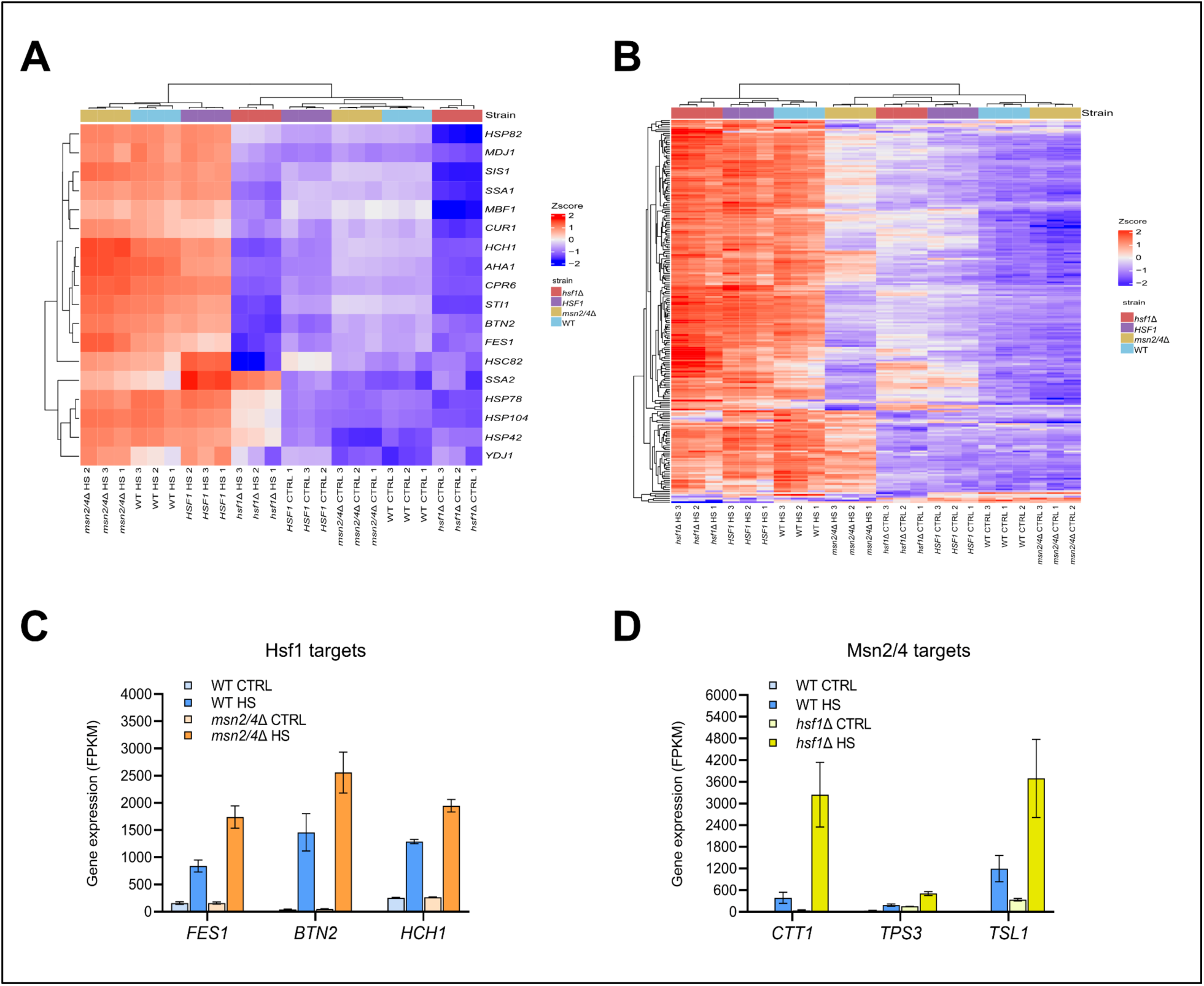
Hsf1 and Msn2/4 display functional compensatory induction following HS. (A) Heat map shows changes in the expression of 18 Hsf1 selective targets upon HS in WT, *hsf1*Δ*, HSF1* and *msn2/4*Δ. (B) Heatmap showing changes in the expression of Msn2/4 targets upon HS in WT, *hsf1*Δ*, HSF1* and *msn2/4*Δ. (C) Gene expression of 3 Hsf1 selective targets in WT and *msn2/4*Δ in untreated cells (CTRL) or upon HS. (D) Gene expression of 3 Msn2/4 targets in WT and *hsf1*Δ upon HS or control conditions (CTRL). Experiment performed in triplicate.

## Discussion

Taken our findings together, we propose the model outlined in Figure 7. Accordingly, Hsf1 is activated under both heat shock and AzC treatment while Msn2/4 are only required for the response to the thermal stress. AzC is incorporated into nascent proteins instead of proline and impairs their folding leading to aggregation. The misfolded proteins sequester Hsp70 and consequently release Hsf1 from its negative regulation resulting in that active Hsf1 binds HSEs and induces the HSR. In contrast, Msn2/4 are not affected by this treatment and remain inactive in the cytoplasm. On the other hand, HS induces broader stress damage that activates both the Hsf1-dependent HSR and the Msn2/4-dependent ESR. Hsf1 and Msn2/4 bind promoters of selective targets as well as of shared targets. In the *hsf1*Δ strain the response to HS is mounted by Msn2/4 and limited to the STRE regulon. Moreover, Hsf1 and Msn2/4 display functional compensatory induction upon HS so that Msn2/4 targets are stronger induced in the *hsf1*Δ strain and, vice versa, Hsf1-selective targets are more highly expressed in the *msn2/4*Δ strain. Thus, Hsf1 and Msn2/4 cooperate together in order to form an effective response to heat stress. In contrast, the response to AzC treatment, hence specifically protein misfolding, only involves Hsf1 and is Msn2/4 independent.

**Figure 7.**
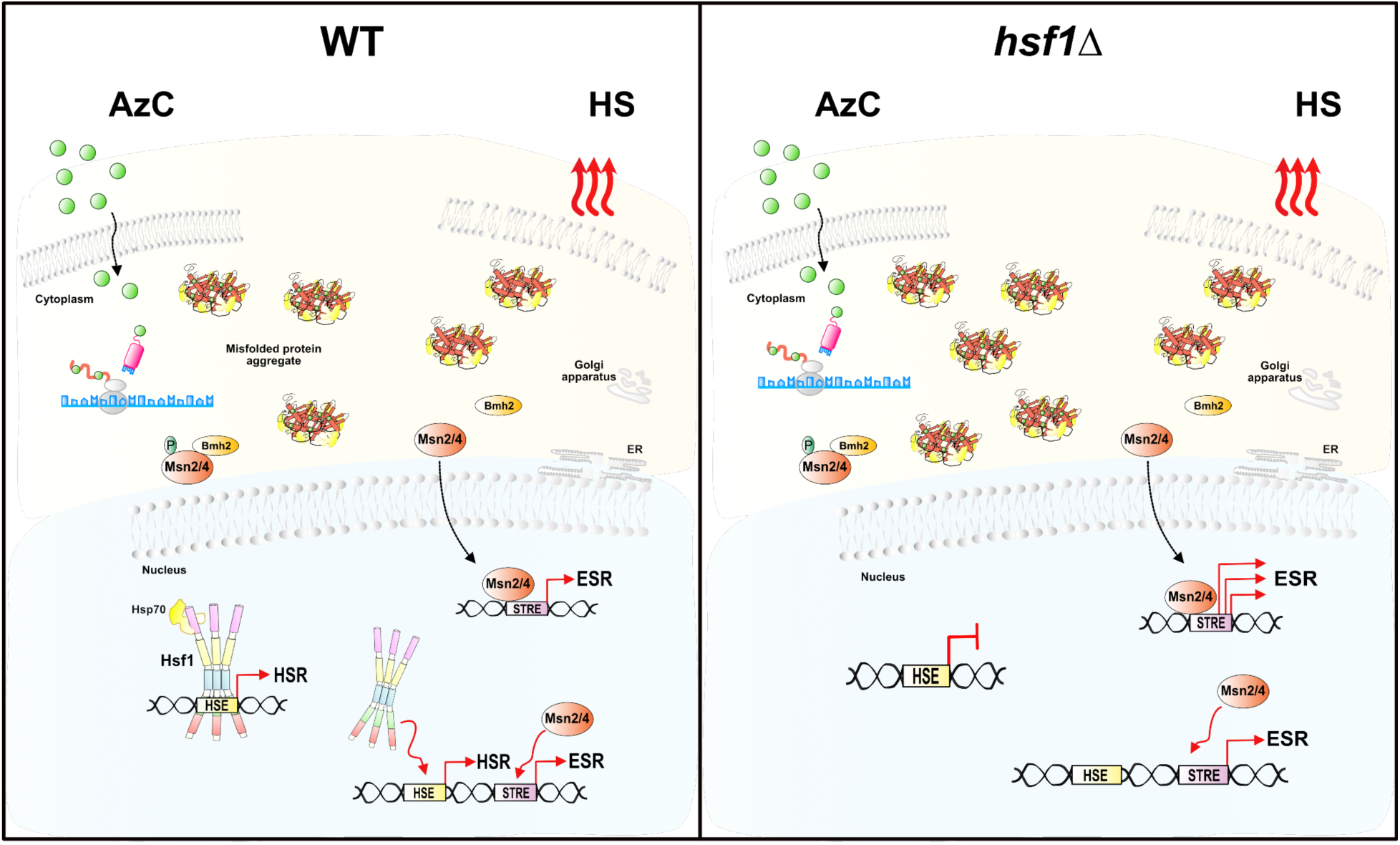
Graphical model showing the response mechanism of *hsf1*Δ to HS and AzC treatment. See Discussion for details

With this work we have for the first time constructed and analyzed a true *hsf1*Δ strain. This strain facilitates the study of *hsf1* phenotypes during both basal and stress conditions and clarifies the specific contribution of Hsf1 to protein homeostasis and stress biology. So far the tools available to study Hsf1 function have been dominant allele *EXA3-1* that results in the expression of Hsf1 with a single amino acid substitution in the DNA binding domain (Nelson *et al*., 1992; Halladay and Craig, 1995) and the transient inactivation of Hsf1 by anchor-away technology (Solís *et al*., 2016). However, the phenotypes of *EXA3-1* strains are somewhat puzzling and include delayed activation of the HSR and an altered induction of Hsf1 targets. Some targets are more strongly induced while others respond more weakly upon HS (Halladay and Craig, 1995). Transient inactivation of Hsf1 using anchor-away technology has been informative. Cells with reduced Hsf1 activity, due to its sequestration outside the nucleus, arrested growth yet provided a transient window for fruitful transcriptome analysis (Solís *et al*., 2016). Inspired by this work that suggested that the Hsp70 and Hsp90 expression was sufficient to replace the essential function of Hsf1, we managed to construct a viable *hsf1*Δ strain. We also found that expression of the Hsp70-activating JDP Ydj1 was required to obtain a viable *hsf1*Δ strain. Analysis of this strain allowed us to conclude that *hsf1*Δ has severely disrupted proteostasis already under basal conditions suggesting that Hsf1 plays a key role in the maintenance of proteostasis even in a cell in which Hsp70 and Hsp90 are constitutively expressed. Hsf1 displays negative chemical-genetic interactions with AzC and its overexpression suppresses AzC toxicity (Berg *et al*., 2020). We find that AzC specifically activates Hsf1 and not Msn2/4. It has previously been suggested that AzC activates Hsf1 and Msn2/4, although the latter more weakly (Trotter *et al*., 2002). Our new findings contrast the previous study, specifically in our dataset we find that *HSP12* induction by AzC is fully dependent on Hsf1 with no contribution of Msn2/4. Moreover, treatment of *hsf1Δ* cells with AzC results in no stress transcription. We have ruled out the scenario that the *hsf1Δ* strain does not experience protein misfolding damage when AzC is administered by detecting the induction of protein aggregates in the cells. Furthermore, the *hsf1Δ* strain is sensitive to AzC demonstrating that it causes damage. The detectable protein misfolded damage appears to be largely limited to the cytoplasm/nucleus. We do not see activation of the UPR by AzC that is known to activate this pathway in mammalian cells (Luo and Lee, 2002; Shang and Lehrman, 2004; Roest *et al*., 2018). We consider the possibility that cytoplasmic proteins are more affected by the treatment with AzC resulting in a higher level of damaged proteins in the cytoplasm than in the ER. Alternatively, the ER proteostasis network may have a higher buffering capacity than its cytosolic counterpart. In light of these results, Hsf1 appears to represent the main and most sensitive monitoring mechanism of misfolded proteins in yeast cells that regulates adaptive transcription.

Hsf1 and Msn2/4 stress activation converges on many shared promoters and their interdependency on the individual gene level is often unclear. In this work we find that Hsf1 and Msn2/4 display functional compensatory induction. We do not think that it is chromatin related since they bind different regions, one possible explanation is that cells accumulate more damage under HS when one of the two pathway is not active. Alternatively, it is possible that there is a negative cross-talk between these transcription factors. Hsf1 is indeed needed when cell proliferation is high and protein synthesis rates are maximized, while Msn2/4 are induced by stressors that lead to reduced cell growth associated with respiration and stationary phase (Brauer *et al*., 2008). It is plausible that they negatively regulate each other in order to balance the two responses. Recently, it has been shown that in aneuploid cells with a chronic state of proteotoxicity the constitutive activation of ESR by Msn2/4 impairs the activation of Hsf1. In these cells, repression of Msn2/4 via engineered activation of PKA revealed significant expression of Hsf1-dependent genes and their prolonged induction following HS. Thus, Msn2/4 negatively modulate the Hsf1 response (Kane *et al*., 2021). Another possibility is that the thermal damage is rapidly counteracted by cooperative action of Hsf1 and Msn2/4 and when one of the two pathways is inactivated, the remaining one becomes stronger induced by the damage due to such lack of the negative feedback regulation. Interestingly, in this regard is that our Hsp70 and Hsp90 overexpressing *HSF1* rescue strain has a modified HSR as well as ESR suggestion that negative regulation by these chaperones may act also in the ESR.

The *hsf1*Δ strain may function as an important tool for future genetic studies of the proteostasis system and the vast biology related to protein misfolding. A major obstacle when studying such protein misfolding phenotypes is that they often are confounded or even masked by the activity of compensatory chaperone factors induced by Hsf1. In the *hsf1*Δ strain the stable levels of Ydj1, Ssa2 and Hsc82 are not dependent on Hsf1 and the entire Hsf1-specific regulon is not responsive to altered levels of protein misfolding. Thus, the *hsf1*Δ allele provides a single kanMX-marked locus that by crossing may be introduced in any yeast strain to assess the direct phenotypes of genetic modification of selected proteostasis factors without the confounding feedback regulation of Hsf1. We also do not fail to note the biotechnological utility of a yeast strain that does not change its transcription in response to the accumulation of misfolded proteins. According to our results, Hsf1 is potentially the only transcription factor present in yeast that in a sensitive manner detects the accumulation of misfolded proteins and induce a transcriptional response. The *hsf1*11 allele effectively makes cells blind to the production of misfolded proteins and facilitates the rational genetic engineering of wanted and unwanted chaperone systems in biotechnological applications as well as in scientific experiments.

## Materials and Methods

### Media and yeast strains

Yeast peptone dextrose (YPD) medium, synthetic complete dextrose (SC) medium and synthetic minimal (SD) medium were used and supplemented to support the growth of auxotrophic strains as previously described (Amberg *et al*., 2005).

The yeast strains used in this study are listed in Table 2. The *hsf1*11 strain was constructed by transforming the diploid strain BY4743 to G418^R^ using *Nde*I, *Pml*I, *SacI*I and *Not*I-restricted pJK074 releasing a knock cassette (Brachmann *et al*., 1998). The resulting heterozygous knock was sporulated and NY129 is a meiotic segregant. The cassette for the genomic deletion of *HSF1* and integration of genes driving the constitutive expression of *YDJ1*, *HSC82* and *SSA2* (*hsf1*11::[***P****_ADH1_-YDJ1; kanMX; **P**_TEF1_-HSC82; **P**_TDH3_-SSA2*]) was assembled on plasmid pJK074. First the *Not*I restriction site on pCA807 (Gowda *et al*., 2013) was removed by *Not*I restriction and relegation. Next, ***P****_TEF1_-HSC82* and *P_TDH3_-SSA2* were introduced by yeast homologous recombination between restricted pCA807 and 4 PCR products: 2 kb *kanMX-**P**_TEF1_*, template pYM-N18 (Janke *et al*., 2004), primers 5’-ATGAATAATGCTGCAAATACAGGGACGACCAATGAGTCAAACGTGAGCGTACGCTGCAGGTCGAC-3’, 5’-GAAATTCAAAAGTTTCACCAGCCATCGATGAATTCTCTGTCG-3’; 2.2 kb *HSC82*, template genomic DNA, primers 5’-CGGACGACAGAGAATTCATCGATGGCTGGTGAAACTTTTG-3’, 5’-TTTGGGCATGTACGGGTTACAGCAGTTAATCAACTTCTTCCATCTC-3’; 0.6 kb *P_TDH3_,* template pCA859 (Gowda *et al*., 2016), primers 5’-CACCGAGATGGAAGAAGTTGATTAACTGCTGTAACCCGTACATGC-3’, 5’-CCTAAATCAATACCGACAGCTTTAGACATTTTGTTTGTTTATGTGTG-3’; 2.1 kb *SSA2*, template genomic DNA, primers 5’-TCGAATAAACACACATAAACAAACAAAATGTCTAAAGCTGTCGGTATTG-3’, 5’-CTATTTCTTAGCTCGTTTGGGCAGGCGGTGATCGTTGTACTCTGTTAATCAACTTCTTCGACAG-3’. Intermediate plasmid pJK067 was rescued as a Amp^R^/Kan^R^ *E. coli* from Ura^+^ yeast transformants. Finally, **P**_ADH1_-*YDJ1* was introduced in *Sma*I/*Pac*I restricted pJK067 using yeast homologous recombination with 2 PCR products: 1.5 kb ***P****_ADH1_*, amplified from pCA148 (C.A. lab collection) primers 5’-GAATAATGCTGCAAATACAGGGACGACCAATGAGTCAAACGTGAGCGAAGAAATGATGGTAAATG-3’, 5’-CGTAAAACTTAGTTTCTTTAACCATTCCGCCCGGAATTAATTCAAG-3’; 1.3 kb *YDJ1*, template pCA893 (Holmberg, Gowda and Andréasson, 2014), primers 5’-CTTGAATTAATTCCGGGCGGAATGGTTAAAGAAACTAAGTTTTACG-3’, 5’-CGGGGACGAGGCAAGCTAAACAGATCTGGCGCGCCTTAATTAACCTCATTGAGATGCACATTGAAC-3’. pJK074 was selected as a Amp^R^/Kan^R^ *E. coli* from Ura^+^ yeast transformants. The plasmid was verified by sequencing.

**Table 2.**
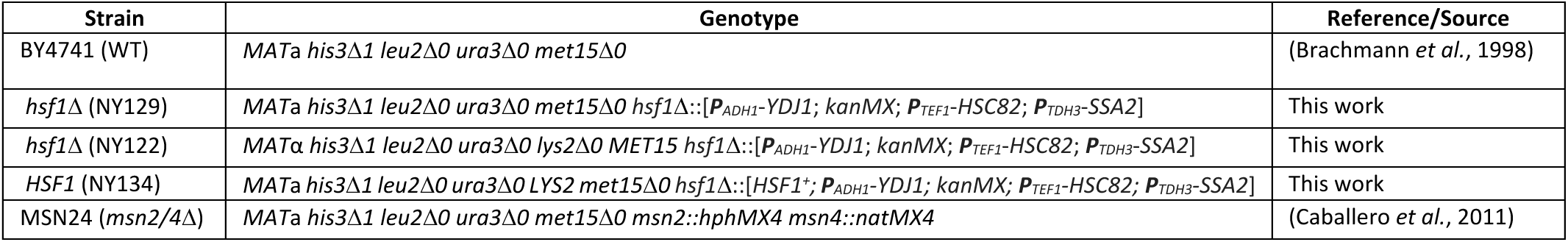
Yeast strains

The *HSF1* rescue strain was obtained by selecting a *hsf1*11 transformant that grew at 37°C after transformation with a 2.7 kb PCR product encompassing the *HSF1* promoter followed by the *HSF1* ORF and sequences homologous to the *ADH1* promoter. The PCR product was obtained from template plasmid pCA807 (Gowda *et al*., 2013) using primers 5’-CTTTGCGAGTGTAAAACTAG-3’; 5’-GTCTTTTGCCTTCATGCTCCTTGATTTCCTATTTCATTTACCATCATTTCTTAGCTCGTTTGGGCAGG-3’.

### Growth assay

Yeast strains were grown overnight in YPD liquid media at 30°C. Next morning, all the strains were diluted to OD_600_= 0.1 (0.2 *hsf1*11) and allowed to grow to logarithmic phase. 1:10 dilution series were performed and cells were spotted onto solid SD medium and were allowed to grow for 3 days at 25°C, 30°C, 37°C and 39°C. For the recovery experiment, 1:10 dilution was performed and cells were heat shocked at 37°C for 15 minutes in a shaking water bath; untreated and heat shocked cells were spotted into SC agar plates that were incubated for 6 days at 30°C. Toxicity of azetidine-2-carboxylic acid (AzC) (AH-diagnostic AB, #SC-263441A) was assessed spotting cell suspensions onto solid YPD medium supplemented with 0, 1, 2, 5 or 10 mM of AzC followed by incubation for 3 days at 25°C or 30°C.

### Isolation of protein aggregates

Protein aggregates were isolated based on a previously published protocol (Jonathan D. Rand and Chris M. Grant, 2006; Masser *et al*., 2019). Briefly, overnight cultures were grown at 25°C or 30°C and was diluted to OD_600_ = 0.1 (0.2 *hsf1*11) in the morning and were grown until OD_600_= 0.8. Cells were treated with 10 mM of AzC for 1h at 25°C and then harvested by centrifugation. Cells were re-suspended in ice-cold lysis buffer (100 mM Tris-HCl pH 7.5, 200 mM NaCl, 1 mM PMSF, 1mM DTT, 5% glycerol) and lysed by 3 rounds in an Emulsiflex B-15 high pressure homogenizer (Avestin, Ottawa, Ontario, Canada) at 25,000 psi. Cell debris was removed by three times repeated centrifugation at 3,000 g for 5 minutes. Protein concentration was set to 1.5 mg/ml using the Bradford assay and 1 ml lysate was used for purification of the aggregates. Lysate was centrifuged at 20,000 g for 2×10 minutes and the pellet was washed with 320 μl of lysis buffer supplemented with 2 % NP-40 and sonicated for 15 minutes with 60% amplitude. Protein aggregates were sedimented by centrifugation at 20,000 g for 2×10 minutes and the pellet re-suspended in 80 μl of 2xSDS sample buffer and boiled for 4 minutes at 95°C. Proteins were separated by SDS-PAGE and visualized by Pierce Silver Staining Kit (Thermo Fisher, #24612).

### Bioluminescent assay for the determination of Hsf1 and Msn2/4 activity

A yNlucPEST reporter was used to monitor Hsf1 and Msn2/4 activity as previously described (Masser A. et al., 2016; Masser *et al*., 2019). Cells were grown at 25°C to logarithmic phase and then heat shocked at 37°C for 15 minutes. 5 μl NanoGlo substrate (Promega GmbH, Germany, #N1110) diluted 1:100 in the supplied lysis buffer and was mixed with the 50 μl culture in a white 96-well plate and bioluminescence was determined after 3 min incubation using an Orion II Microplate Luminometer (Berthold Technologies GmbH and Co. KG, Germany). Bioluminescence light units (BLU) are defined as the relative light units (RLU)/s of 1 mL cells at OD600 = 1.0.

### In vivo firefly luciferase reactivation assay

In vivo firefly luciferase reactivation was performed as previously described (Abrams and Morano, 2013; Kaimal *et al*., 2017). Briefly, cells carrying firefly luciferase reporter (FFL-GFP-NES) were grown to OD_600_= 1 at 30°C and then treated with 100 mg/liter of cycloheximide (CHX) or CHX and 4 mM of guanidine hydrochloride (GnHCl) together. After treatment cells were heat shocked at 43°C for 15 min in a water bath. The FFL-GFP reactivation at 30°C was monitored by using Orion II Microplate Luminometer with 100 μl culture mixed with 50 μl of D-luciferin 455 μg/ml.

### Protein extraction, SDS-PAGE and immunoblot analysis

Protein extracts were prepared as previously described (Silve *et al*., 1991; Gowda *et al*., 2013). Briefly, cell culture aliquots were incubated on ice for 10 minutes after addition of precooled NaOH to a final concentration of 0.37 M. Trichloroacetic acid (TCA) was then added to a final concentration of 8.3% (w/v). The day after, cells were collected by centrifugation and the pellet washed in 1 M Tris base. The final pellet was re-suspended in 2xSDS-sample buffer and boiled at 95°C for 5 minutes. Equal amounts of SDS-solubilized proteins were separated using SDS-PAGE and transferred to a nitrocellulose membranes. Membranes were incubated for 1 h in 5% nonfat dry milk in TBST (Tris-buffered saline/0.1% Tween) and then incubated over night with the following primary antibodies: α-Hsp104 (rabbit 1:5000, Enzo ADI-SPA-1040), α-Hsp42 (rabbit 1:5000), α-Fes1 (rabbit 1:5000), α-Hsc82/Hsp82 (rabbit 1:50000), α-Ssa1/2 (rabbit 1:67000), α-Ydj1 (1:4000 mouse, Sigma-Aldrich #SAB5200007), α-Sse1 (rabbit 1:100000) and α-GFP (rabbit 1:5000, Invitrogen #A6455). α-Pgk1 (mouse 1:5000, ThermoFisher, #22C5D8) and α-GAPDH (mouse 1:5000, Invitrogen #MA5-15738), were used as a loading control. The day after membranes were washed 3 times with TBST and then incubated for 1 h at room temperature with secondary antibodies. Proteins were visualized using the Odyssey Fc infrared imaging system (Li-COR Biosciences). Signal quantification was performed using the Image Studio Lite software (LICOR Biosciences) by normalizing the background-corrected signals to the loading control.

### mRNA sequencing (mRNA-Seq)

Cells were grown in triplicates for each condition in YPD liquid media at 25°C until OD_600_ = 0.8. Cells were heat shocked at 37°C for 15 min in a shaking water bath or treated with 10 mM AzC for 1h at 25°C or left untreated before harvesting them by centrifugation. RNA was extracted using RiboPure Yeast RNA Purification Kit (Thermo Fisher Scientific, #AM1926). The quality of the RNA extracted was analyzed using high sensitivity RNA BioAnalyzer Chip (Agilent). The RNA-seq was performed by National Genomics Infrastructure Stockholm Node, Science for Life Laboratory. Libraries were prepared using Illumina TruSeq RNA poly-A selection prep and the sequencing was performed on Illumina NovaSeq6000 SP flowcell. Reference Genome: Budding yeast (Saccharomyces cerevisiae, R64-1-1). FPKM were generated using the bioinformatics pipeline nf-core/rnaseq pipeline. Principal component analysis was performed based on gene FPKMs using plotPCA function in R (v 4.2.1). Differential gene expression analysis was performed using DESeq2 (v 1.36.0). Heatmaps were generated using the ComplexHeatmap package with scaled, variance stabilized RNA abundances (Gu *et al*., 2016). Genes with adjusted p value < 0.01 and absolute log2 Fold Change > 1 are considered significant.

### Statistical analysis

Error bars show standard deviation of at least three biological independent replicates (starting from independent cell cultures). Statistical testing was done with Graphpad Prism 9 with one-way ANOVA multiple comparisons. P-values are defined as *p<0.05, **p<0.01 and ***p<0.001.

## Acknowledgments

We thank Thomas Nyström for supplying the *msn2/4*Δ strain. We acknowledge the National Genomics Infrastructure (NGI) at SciLifeLab for RNA-seq. This work was supported by Knut and Alice Wallenberg Foundation, Swedish Research Council (project grant 2019-04052 to CA) and the Swedish Cancer Society (20 1045 to CA).

## Notes

### Competing Interest Statement

The authors have declared no competing interest.

## References

Abrams, JL, and Morano, KA (2013). Coupled assays for monitoring protein refolding in Saccharomyces cerevisiae. J Vis Exp, 2–7.

Amberg, DC, Burke, DJ, and Strathern, JN (2005). Methods in Yeast Genetics: A Cold Spring Harbor Laboratory Course Manual, 2005 Edition. A Cold Spring Harb Lab Course Man 5, 205.

Beck, T, and Hall, MN (1999). The TOR signalling pathway controls nuclear localization of nutrient-regulated transcription factors. Nature 402, 689–692.

Berg, MD, Zhu, Y, Isaacson, J, Genereaux, J, Loll-Krippleber, R, Brown, GW, and Brandl, CJ (2020). Chemical-genetic interactions with the proline analog l-azetidine-2-carboxylic acid in saccharomyces cerevisiae. G3 Genes, Genomes, Genet 10, 4335–4345.

Bonner, JJ, Ballou, C, and Fackenthal, DL (1994). Interactions between DNA-bound trimers of the yeast heat shock factor. Mol Cell Biol 14, 501–508.

Bonner, JJ, Carlson, T, Fackenthal, DL, Paddock, D, Storey, K, and Lea, K (2000). Complex regulation of the yeast heat shock transcription factor. Mol Biol Cell 11, 1739–1751.

Boy-Marcotte, E, Lagniel, G, Perrot, M, Bussereau, F, Boudsocq, A, Jacquet, M, and Labarre, J (1999). The heat shock response in yeast: Differential regulations and contributions of the Msn2p/Msn4p and Hsf1p regulons. Mol Microbiol 33, 274–283.

Brachmann, CB, Davies, A, Cost, GJ, Caputo, E, Li, J, Hieter, P, and Boeke, JD (1998). Designer deletion strains derived from Saccharomyces cerevisiae S288C: A useful set of strains and plasmids for PCR-mediated gene disruption and other applications. Yeast 14, 115–132.

Brauer, MJ, Curtis Huttenhower, EMA, Rosenstein, R, John C. Matese, DG, Boer, VM, Troyanskaya, OG, and Botstein, D (2008). Coordination of Growth Rate, Cell Cycle, Stress Response, and Metabolic Activity in Yeast. Mol Biol Cell 18, 3250–3263.

Caballero, A, Ugidos, A, Liu, B, Öling, D, Kvint, K, Hao, X, Mignat, C, Nachin, L, Molin, M, and Nyström, T (2011). Absence of Mitochondrial Translation Control Proteins Extends Life Span by Activating Sirtuin-Dependent Silencing. Mol Cell 42, 390–400.

Chapman, RE, and Walter, P (1997). Translational attenuation mediated by an mRNA intron. Curr Biol 7, 850–859.

Cox, JS, and Walter, P (1996). A novel mechanism for regulating activity of a transcription factor that controls the unfolded protein response. Cell 87, 391–404.

Fowden, L., and Richmond, MH (1963) (1963). Replacement of proline by azetid|ne-2-carboxylic acid. 71, 1–9.

Gasch, AP, Spellman, PT, Kao, CM, Carmel-Harel, O, Eisen, MB, Storz, G, Botstein, D, and Brown, PO (2000). Genomic expression programs in the response of yeast cells to environmental changes. Mol Biol Cell 11, 4241–4257.

Glover, JR, and Lindquist, S (1998). Hsp104, Hsp70, and Hsp40: A novel chaperone system that rescues previously aggregated proteins. Cell 94, 73–82.

Görner, W, Durchschlag, E, Martinez-Pastor, MT, Estruch, F, Ammerer, G, Hamilton, B, Ruis, H, and Schüller, C (1998). Nuclear localization of the C2H2 zinc finger protein Msn2p is regulated by stress and protein kinase A activity. Genes Dev 12, 586–597.

Gowda, NKC, Kaimal, JM, Masser, AE, Kang, W, Friedländer, MR, and Andréasson, C (2016). Cytosolic splice isoform of Hsp70 nucleotide exchange factor Fes1 is required for the degradation of misfolded proteins in yeast. Mol Biol Cell 27, 1210–1219.

Gowda, NKC, Kandasamy, G, Froehlich, MS, Jürgen Dohmen, R, and Andréasson, C (2013). Hsp70 nucleotide exchange factor Fes1 is essential for ubiquitin-dependent degradation of misfolded cytosolic proteins. Proc Natl Acad Sci U S A 110, 5975–5980.

Grably, MR, Stanhill, A, Tell, O, and Engelberg, D (2002). HSF and Msn2/4p can exclusively or cooperatively activate the yeast HSP104 gene. Mol Microbiol 44, 21–35.

Gross, DS, Adams, CC, Lee, S, and Stentz, B (1993). A critical role for heat shock transcription factor in establishing a nucleosome-free region over the TATA-initiation site of the yeast HSP82 heat shock gene. EMBO J 12, 3931– 3945.

Gross, DS, English, KE, Collins, KW, and Lee, S (1990). Genomic footprinting of the yeast HSP82 promoter reveals marked distortion of the DNA helix and constitutive occupancy of heat shock and TATA elements. J Mol Biol 216, 611–631.

Gu, Z, Eils, R, and Schlesner, M (2016). Complex heatmaps reveal patterns and correlations in multidimensional genomic data. Bioinformatics 32, 2847–2849.

Hahn, J-S, Hu, Z, Thiele, DJ, and Iyer, VR (2004). Genome-Wide Analysis of the Biology of Stress Responses through Heat Shock Transcription Factor. Mol Cell Biol 24, 5249–5256.

Hall, MP et al. (2012). Engineered luciferase reporter from a deep sea shrimp utilizing a novel imidazopyrazinone substrate. ACS Chem Biol 7, 1848–1857.

Halladay, JT, and Craig, EA (1995). A heat shock transcription factor with reduced activity suppresses a yeast HSP70 mutant. Mol Cell Biol 15, 4890–4897.

Holmberg, MA, Gowda, NKC, and Andréasson, C (2014). A versatile bacterial expression vector designed for single-step cloning of multiple DNA fragments using homologous recombination. Protein Expr Purif 98, 38–45.

Jakobsen, BK, and Pelham, HR (1988). Constitutive binding of yeast heat shock factor to DNA in vivo. Mol Cell Biol 8, 5040–5042.

Janke, C et al. (2004). A versatile toolbox for PCR-based tagging of yeast genes: New fluorescent proteins, more markers and promoter substitution cassettes. Yeast 21, 947–962.

Jonathan D. Rand and Chris M. Grant (2006). The Thioredoxin System Protects Ribosomes against Stress-induced Aggregation. Mol Biol Cell 16, 5356–5372.

Jung, G, and Masison, DC (2001). Guanidine hydrochloride inhibits Hsp104 activity in vivo: A possible explanation for its effect in curing yeast prions. Curr Microbiol 43, 7–10.

Kaimal, JM, Kandasamy, G, Gasser, F, and Andréasson, C (2017). Coordinated Hsp110 and Hsp104 Activities Power Protein Disaggregation in Saccharomyces cerevisiae. Mol Cell Biol 37.

Kampinga, HH, Andreasson, C, Barducci, A, Cheetham, ME, and Cyr, D (2019). Function, evolution, and structure of J-domain proteins. 7–15.

Kane, AJ, Brennan, CM, Xu, AE, Solís, EJ, Terhorst, A, Denic, V, and Amon, A (2021). Cell adaptation to aneuploidy by the environmental stress response dampens induction of the cytosolic unfolded-protein response. Mol Biol Cell 32, 1557–1564.

Kawahara, T, Yanagi, H, Yura, T, and Mori, K (1997). Endoplasmic reticulum stress-induced mRNA splicing permits synthesis of transcription factor Hac1p/Ern4p that activates the unfolded protein response. Mol Biol Cell 8, 1845–1862.

Keefer, KM, and True, HL (2017). A toxic imbalance of Hsp70s in Saccharomyces cerevisiae is caused by competition for cofactors. Mol Microbiol 105, 860–868.

Kim, I-S, Kim, H, Kim, Y-S, Jin, I, and Yoon, H-S (2013). HSF1-mediated oxidative stress response to menadione in Saccharomyces cerevisiae KNU5377Y3 by using proteomic approach. Adv Biosci Biotechnol 04, 44–54.

Krakowiak, J, Zheng, X, Patel, N, Feder, ZA, Anandhakumar, J, Valerius, K, Gross, DS, Khalil, AS, and Pincus, D (2018). Hsf1 and Hsp70 constitute a two-component feedback loop that regulates the yeast heat shock response. Elife 7.

Kuang, Z, Pinglay, S, Ji, H, and Boeke, JD (2017). Msn2/4 regulate expression of glycolytic enzymes and control transition from quiescence to growth. Elife 6, 1–16.

Lambowitz, AM, Kobayashi, GS, Painter, A, and Medoff, G (1983). fT. 25–27.

Li, N, Zhang, LM, Zhang, KQ, Deng, JS, Prändl, R, and Schöffl, F (2006). Effects of heat stress on yeast heat shock factor-promoter binding in vivo. Acta Biochim Biophys Sin (Shanghai) 38, 356–362.

Liu, Y, and Chang, A (2008). Heat shock response relieves ER stress. EMBO J 27, 1049–1059.

Luo, S, and Lee, AS (2002). Requirement of the p38 mitogen-activated protein kinase signalling pathway for the induction of the 78 kDa glucose-regulated protein/immunoglobulin heavy-chain binding protein by azetidine stress: Activating transcription factor 6 as a target for stress-i. Biochem J 366, 787–795.

Martínez-Pastor, MT, Marchler, G, Schüller, C, Marchler-Bauer, A, Ruis, H, and Estruch, F (1996). The Saccharomyces cerevisiae zinc finger proteins Msn2p and Msn4p are required for transcriptional induction through the stress-response element (STRE). EMBO J 15, 2227–2235.

Masser A. et al., 2016 (2016). Identi cation of an NADH-dependent 5-hydroxymethylfurfural-reducing alcohol dehydrogenase in. Yeast, 191–198.

Masser, AE, Ciccarelli, M, and Andréasson, C (2020). Hsf1 on a leash – controlling the heat shock response by chaperone titration. Exp Cell Res 396, 112246.

Masser, AE, Kang, W, Roy, J, Kaimal, JM, Quintana-Cordero, J, Friedländer, MR, and Andréasson, C (2019). Cytoplasmic protein misfolding titrates Hsp70 to activate nuclear Hsf1. Elife 8, 1–27.

McDaniel, D, Caplan, AJ, Lee, MS, Adams, CC, Fishel, BR, Gross, DS, and Garrard, WT (1989). Basal-level expression of the yeast HSP82 gene requires a heat shock regulatory element. Mol Cell Biol 9, 4789–4798.

Morano, KA, Grant, CM, and Moye-Rowley, WS (2012). The response to heat shock and oxidative stress in saccharomyces cerevisiae. Genetics 190, 1157–1195.

Morano, KA, Santoro, N, Koch, KA, and Thiele, DJ (1999). A trans -Activation Domain in Yeast Heat Shock Transcription Factor Is Essential for Cell Cycle Progression during Stress. Mol Cell Biol 19, 402–411.

Mori, K, Kawahara, T, Yoshida, H, Yanagi, H, and Yura, T (1996). Signalling from endoplasmic reticulum to nucleus: Transcription factor with a basic-leucine zipper motif is required for the unfolded protein-response pathway. Genes to Cells 1, 803–817.

Nelson, RJ, Heschl, MFP, and Craig, EA (1992). Isolation and characterization of extragenic suppressors of mutations in the SSA hsp70 genes of Saccharomyces cerevisiae. Genetics 131, 277–285.

Parsell, DA, Kowal, AS, Singer, MA, and Lindquist, S (1994). Protein disaggregation. Nature 372, 475–478.

Patriarcaandbrunomaresca, EJ (1990). Thermotolerance following Heat Shock Prevents impairment of Mitochondrial A levated Temperatures in Saccharom. 64, 57–64.

Peter Lee et al., 2008 (2008). YAK1 Msn24.pdf.

Pincus, D, Anandhakumar, J, Thiru, P, Guertin, MJ, Erkine, AM, and Gross, DS (2018). Genetic and epigenetic determinants establish a continuum of Hsf1 occupancy and activity across the yeast genome. Mol Biol Cell 29, 3168–3182.

Ritossa, F (1996). Discovery of the heat shock response. Cell Stress Chaperones 1, 97–98.

Roest, G, Hesemans, E, Welkenhuyzen, K, Luyten, T, Engedal, N, Bultynck, G, and Parys, JB (2018). The ER stress inducer l-azetidine-2-carboxylic acid elevates the levels of phospho-eiF2α and of LC3-II in a ca2+-dependent manner. Cells 7.

Ron, D, and Walter, P (2007). Signal integration in the endoplasmic reticulum unfolded protein response. Nat Rev Mol Cell Biol 8, 519–529.

Sakurai, H, and Takemori, Y (2007). Interaction between heat shock transcription factors (HSFs) and divergent binding sequences: Binding specificities of yeast HSFs and human HSF1. J Biol Chem 282, 13334–13341.

Santoro, N, Johansson, N, and Thiele, DJ (1998). Heat Shock Element Architecture Is an Important Determinant in the Temperature and Transactivation Domain Requirements for Heat Shock Transcription Factor. Mol Cell Biol 18, 6340–6352.

Sathyanarayanan, U, Musa, M, Bou Dib, P, Raimundo, N, Milosevic, I, and Krisko, A (2020). ATP hydrolysis by yeast Hsp104 determines protein aggregate dissolution and size in vivo. Nat Commun 11.

Schmitt, AP, and Mcentee, K (1996). Msn2p, a zinc finger DNA-binding protein, is the transcriptional activator of the multistress response in Saccharomyces cerevisiae. Proc Natl Acad Sci U S A 93, 5777–5782.

Shang, J, and Lehrman, MA (2004). Discordance of UPR signaling by ATF6 and Ire1p-XBP1 with levels of target transcripts. Biochem Biophys Res Commun 317, 390–396.

Sidrauski, C, and Walter, P (1997). The transmembrane kinase Ire1p is a site-specific endonuclease that initiates mRNA splicing in the unfolded protein response. Cell 90, 1031–1039.

Silve, S, Volland, C, Garnier, C, Jund, R, Chevallier, MR, and Haguenauer-Tsapis, R (1991). Membrane insertion of uracil permease, a polytopic yeast plasma membrane protein. Mol Cell Biol 11, 1114–1124.

Solís, EJ, Pandey, JP, Zheng, X, Jin, DX, Gupta, PB, Airoldi, EM, Pincus, D, and Denic, V (2016). Defining the Essential Function of Yeast Hsf1 Reveals a Compact Transcriptional Program for Maintaining Eukaryotic Proteostasis. Mol Cell 63, 60–71.

Sorger, PK, and Pelham, HR (1987a). Purification and characterization of a heat-shock element binding protein from yeast. EMBO J 6, 3035–3041.

Sorger, PK, and Pelham, HR (1987b). Purification and characterization of a heat-shock element binding protein from yeast. EMBO J 6, 3035–3041.

Steurer, C, Eder, N, Kerschbaum, S, Wegrostek, C, Gabriel, S, Pardo, N, Ortner, V, Czerny, T, and Riegel, E (2018). HSF1 mediated stress response of heavy metals. PLoS One 13, 1–18.

Toivola, DM, Strnad, P, Habtezion, A, and Omary, MB (2010). Intermediate filaments take the heat as stress proteins. Trends Cell Biol 20, 79–91.

Treger, JM, Schmitt, AP, Simon, JR, and McEntee, K (1998). Transcriptional factor mutations reveal regulatory complexities of heat shock and newly identified stress genes in Saccharomyces cerevisiae. J Biol Chem 273, 26875–26879.

Trotter, EW, Kao, CMF, Berenfeld, L, Botstein, D, Petsko, GA, and Gray, J V. (2002). Misfolded proteins are competent to mediate a subset of the responses to heat shock in Saccharomyces cerevisiae. J Biol Chem 277, 44817– 44825.

Verghese, J, Abrams, J, Wang, Y, and Morano, KA (2012). Biology of the Heat Shock Response and Protein Chaperones: Budding Yeast (Saccharomyces cerevisiae) as a Model System. Microbiol Mol Biol Rev 76, 115– 158.

Voellmy and Boellmann, 2007 (2007). Chaperone Regulation of the Heat Shock Protein Response.

Vogel, JL, Parsell, DA, and Lindquist, S (1995). Heat-shock proteins Hsp104 and Hsp70 reactivate mRNA splicing after heat inactivation. Curr Biol 5, 306–317.

Wang, H-Y, Fu, JC-M, Lee, Y-C, and Lu, P-J (2013). Hyperthermia Stress Activates Heat Shock Protein Expression via Propyl Isomerase 1 Regulation with Heat Shock Factor 1. Mol Cell Biol 33, 4889–4899.

Welch, WI, and Suhan, JP (1985). Morphological study of the mammalian stress response: Characterization of changes in cytoplasmic organelles, cytoskeleton, and nucleoli, and appearance of intranuclear actin filaments in rat fibroblasts after heat-shock treatment. J Cell Biol 101, 1198–1211.

De Wever, V, Reiter, W, Ballarini, A, Ammerer, G, and Brocard, C (2005). A dual role for PP1 in shaping the Msn2-dependent transcriptional response to glucose starvation. EMBO J 24, 4115–4123.

Wiederrecht, G, Seto, D, and Parker, CS (1988). Isolation of the gene encoding the S. cerevisiae heat shock transcription factor. Cell 54, 841–853.

Yamamoto, A, Mizukami, Y, and Sakurai, H (2005). Identification of a novel class of target genes and a novel type of binding sequence of heat shock transcription factor in Saccharomyces cerevisiae. J Biol Chem 280, 11911– 11919.

Zagari, A, Némethy, G, and Heraga, HA (1990). The effect of the L-azetidine-2-carboxylic acid residue on protein conformation. II. Homopolymers and copolymers. Biopolymers 30, 961–966.

Zagari, A, Palmer, KA, Gibson, KD, Némethy, G, and Scheraga, HA (1994). The effect of the L-azetidine-2-carboxylic acid residue on protein conformation. IV. Local substitutions in the collagen triple helix. Biopolymers 34, 51–60.

Zheng, X, Krakowiak, J, Patel, N, Beyzavi, A, Ezike, J, Khalil, AS, and Pincus, D (2016). Dynamic control of Hsf1 during heat shock by a chaperone switch and phosphorylation. Elife 5, 1–26.

